# Peripheral Inflammation is accompanied by Cerebral Hypoperfusion in Mice

**DOI:** 10.1101/2024.09.03.610646

**Authors:** Afolashade Kazeem, Chuang Ge, Maral Tajerian

**Author notes:** Corresponding author: Dr. Maral Tajerian: Biology department, Natural Sciences Building Unit E-104. 65-30 Kissena Blvd, Flushing, NY, 11367.

## Abstract

**Introduction:** Chronic pain is a disabling condition that is accompanied by neuropsychiatric comorbidities such as anxiety, depression, and cognitive decline. While the peripheral alterations are well-studied, we lack an understanding of how these peripheral changes can result in long-lasting brain alterations and the ensuing behavioral phenotypes. This study aims to quantify changes in cerebral blood perfusion using laser speckle contrast imaging (LSCI) in the murine Complete Freund’s adjuvant (CFA) model of unilateral peripheral inflammation.

**Methods:** Twenty female and 24 male adult C57BL/6 mice were randomly assigned to control (0.05ml saline) or 1 of 3 experimental groups receiving CFA (0.01ml, 0.05ml, and 0.1ml) on the right hindpaw. Three days after the intraplantar injections, animals were assessed for signs of pain (von Frey), working memory (y-maze), and anxiety (zero maze and open field), and subjected to craniotomy and *in vivo* LSCI of the parietal-temporal lobes.

**Results:** Unilateral administration of CFA resulted in signs of local inflammation, decreased mechanical thresholds on the affected hindpaw, signs of anxiety in the zero maze, as well as cerebral hypoperfusion in dose-dependent manner.

**Discussion:** To our knowledge, this is the first study using laser speckle contrast imaging to examine the effects of CFA-induced peripheral inflammation on cerebral blood perfusion. It serves as a first step in delineating the path by which insult to peripheral tissues can cause long-lasting brain plasticity via vascular mechanisms.

## Introduction

Inflammation, an adaptive defensive response under homeostatic conditions, plays a crucial role in driving tissue repair and fighting likely pathogens. Pain is one of the cardinal signs of inflammation potentially caused by released mediators forming an “inflammatory soup” capable of nociceptor sensitization [13]. Under pathological conditions, inflammation could lead to chronic pain often presenting with multiple co-morbid psychiatric disorders, including mood alterations [22] and cognitive impairment [6]. One pivotal mechanism that could explain the chronification of pain as well as its resistance to classical treatment is the concept of pain centralization, where initial sensory events can gradually alter the central nervous system, resulting in amplified pain and/or aberrant pain that exists without peripheral sensitization. Alterations in brain circuitry have been extensively reported across a spectrum of pain conditions, such as complex regional pain syndrome [17; 26; 31-33], fibromyalgia [18; 23], neuropathic pain [2; 3; 5; 10; 11; 14; 30], and migraine [7], thus prompting the quest for treatments that could reset these systems. However, most of the research in this area overlooks the evolving brain microvascular changes that may parallel cellular plasticity.

The expensive and somewhat primitive nature of minimally-invasive techniques that could be used to study the pain brain in animal models have historically posed significant practical limitations. The rapid technological advances in the field of *in vivo* blood perfusion imaging paired with the high clinical relevance of such studies have resulted in renewed interest in the role of brain vascular changes and its role in linking peripheral insults to central nervous system (CNS) alterations [35], with laser speckle imaging gaining popularity as a method of *in vivo* perfusion quantitation in the clinic [15; 24].

This study aims to quantify changes in mechanical sensitivity, cognitive function, anxiety, as well as cerebral blood perfusion using laser speckle contrast imaging (LSCI) in the murine Complete Freund’s adjuvant (CFA) model of unilateral peripheral inflammation. This effort is a first step in addressing the identity of the “black box” between central and peripheral mechanism of pain, thereby opening the door to entirely novel therapeutic venues that do not only target the injured tissues but rather address the node of pain chronification.

## Materials and methods

### Animals

Twenty female and 24 male C57BL/6, 12-16 weeks of age, were purchased from a commercial supplier (Jackson lab, USA) and habituated for 14 days at the Queens College animal facility before the start of the experiment. Animals were housed in groups of 3/cage on a 12-hour light/dark cycle and an ambient temperature of 20°C to 22°C, with food and water available *ad libitum*. All animal procedures were approved by the Queens College Institutional Animal Care and Use Committee (Flushing, NY, USA) and conform to the NIH guidelines [1] and the “animal subjects” guidelines of the International Association for the Study of Pain.

### Induction of peripheral inflammation

Animals were randomly assigned to control or one of three experimental groups as described below. Normal saline or undiluted CFA (Sigma-Aldrich, Munich, Germany) were administered to the intraplantar surface of the right hindpaw under isoflurane anesthesia. Signs of hindpaw swelling and abnormal gait were noted by the observer as qualitative measures 3 days after the unilateral injections.

<u>a. Control group:</u> 0.05ml of normal saline; n=11. <u>b. Experimental groups:</u> Group 1: 0.01ml CFA; n=11. Group 2: 0.05ml CFA; n=11. Group 3: 0.1ml CFA; n=11.

### Behavioral testing

All analyses were blinded to the identity and experimental condition of the animal. Mice were habituated to the testing room for 1hr prior to the start of the experiments. All behavioral apparatuses were cleaned and deodorized using 0.325% acetic acid (v/v). All videos were recorded using GoPro cameras and analyzed using the automatic animal tracking software Behaviorcloud©.

The following groups were used: <u>a. Control group:</u> 0.05ml of normal saline; n=5. <u>b. Experimental groups:</u> Group 1: 0.01ml CFA; n=5. Group 2: 0.05ml CFA; n=5. Group 3: 0.1ml CFA; n=5.

#### Mechanical sensitivity

Calibrated monofilaments (Stoelting Co., USA) were applied to the plantar surface of the hind paw and the 50% threshold to withdraw (grams) was calculated as previously described [12]. The stimulus intensity ranged from 0.004 to 1.7g, corresponding to filament numbers 1.65, 2.36, 2.44, 2.83, 3.22, 3.61, 3.84, 4.08, 4.17, and 4.31. For each animal, the actual filaments used within the aforementioned series were determined based on the lowest filament to evoke a positive response (response = flexion reflex) followed by five consecutive stimulations using the up–down method. The filament range and average interval were then incorporated along with the response pattern into each individual threshold calculation.

#### Y-maze

Rodents’ natural inclination to explore new environments was evaluated by quantifying spontaneous alternation in exploring the arms of the maze. In a Y-shaped maze with three identical arms (A, B, and C), mice tend to choose a new arm over one they’ve already visited. The maze, was made in-house using opaque white acrylic. Mice were placed in the center and allowed to explore for 5 minutes. Mice with strong spatial working memory typically entered a new arm without revisiting a previous one. The test measured spontaneous alternation by tracking unique sequences of arm entries (e.g., ABC, BCA). The percentage of alternation was calculated by dividing the number of unique sequences by the total arm entries minus two.

#### Zero maze

The apparatus was made in-house and had the following dimensions: inner circle diameter=47 cm, outer circle diameter=54 cm, height =25cm. Mice were placed at the boundary between the open and enclosed regions and allowed to explore freely for six minutes. The time spent by the mice in each of these regions was quantified.

#### Open field

The apparatus was built in-house as a 28×28×28 (LXWXH) acrylic cube. Mice were placed in the center of the field and allowed to explore for 6 minutes. The time spent in the central 25% area vs the entire arena was calculated in addition to the speed of locomotion in the central and peripheral areas.

### Craniotomy

Mice were weighed and dexamethasone (5ml/mg, i.p.) was administered to prevent/reduce the occurrence of cerebral edema caused by the side effects of isoflurane and the impact of drilling the skull. Under isoflurane anesthesia, mice were transferred to a stereotaxic frame and positioned on a heating pad with continuous temperature monitoring, with sterile ointment applied to the eyes to prevent dryness. The scalp was shaved and cleaned with povidone before incision and drilling (drill bit diameter=0.8mm). The mouse was monitored continuously during surgery, the drill bit was cooled intermittently, and the bone and tissues were moistened with normal saline to prevent dryness. A skull window was created, exposing the cerebrum’s left and right parietal-temporal lobes.

### Laser speckle contrast imaging

The following groups were used: <u>a. Control group:</u> 0.05ml of normal saline; n=6. <u>b. Experimental groups:</u> Group 1: 0.01ml CFA; n=6. Group 2: 0.05ml CFA; n=6. Group 3: 0.1ml CFA; n=6.

The Laser speckle imaging system was used to capture and analyze blood flow dynamics [30] in the brain tissue 3 days post-CFA/saline administration. The setup includes a coherent laser light source, a camera for capturing speckle patterns, and RFLSI iii computer software for data analysis (RWD Life Sciences, Shenzhen, China). Following the skull window establishment, the mouse brain tissue was exposed to laser intensity of 80mw, and white light intensity was level 4 for a recording duration of 12s. The exposure time was 8ms, and the HD temporal algorithm was 2048*2048. Blood perfusion data was analyzed for a constant regions across all animals (0.7cm X 0.5cm) using the RFLSI iii software.

### Statistical Analysis

Data analysis was carried out using one-way analysis of variance (ANOVA) followed by Dunnett’s post hoc test for multiple comparisons. Significance was set at P value <0.05; GraphPad Prism V8.0 (GraphPad Software, San Diego, USA).

## Results

Unilateral injection of CFA into the mouse hindpaw resulted in classic signs of inflammation per the subjective observation of the experimenter 3 days after treatment. The administration of saline was not linked to signs of hindpaw swelling or abnormal gait. Meanwhile, the low dose of 0.01mL CFA caused mild swelling in the right paw and abnormal gait, the medium dose of 0.05mL CFA resulted in swelling extending beyond the paw as well as abnormal gait, and the high dose of 0.1mL caused swelling extending farther beyond the paw and abnormal gait.

To provide a less subjective assessment of behavioral changes related to paw inflammation, we conducted tests measuring mechanical sensitivity (von Frey), working memory (y-maze), anxiety-like behaviors (zero maze, open field), and overall locomotor function (open field). Our results indicate a significant increase in mechanical sensitivity following 0.1ml CFA administration (**Figure 1A**), as shown by the difference in thresholds between the unaffected and injured paws [ΔvF threshold (g)= threshold_contra_ (g)-threshold_ipsi_ (g); one-way ANOVA p=0.02, F=4.358, Dunnett’s multiple comparison test with an adjusted P value of 0.007, mean difference between the control (0ml CFA) and high dose (0.1ml CFA) = -0.7323].

**Figure 1:**
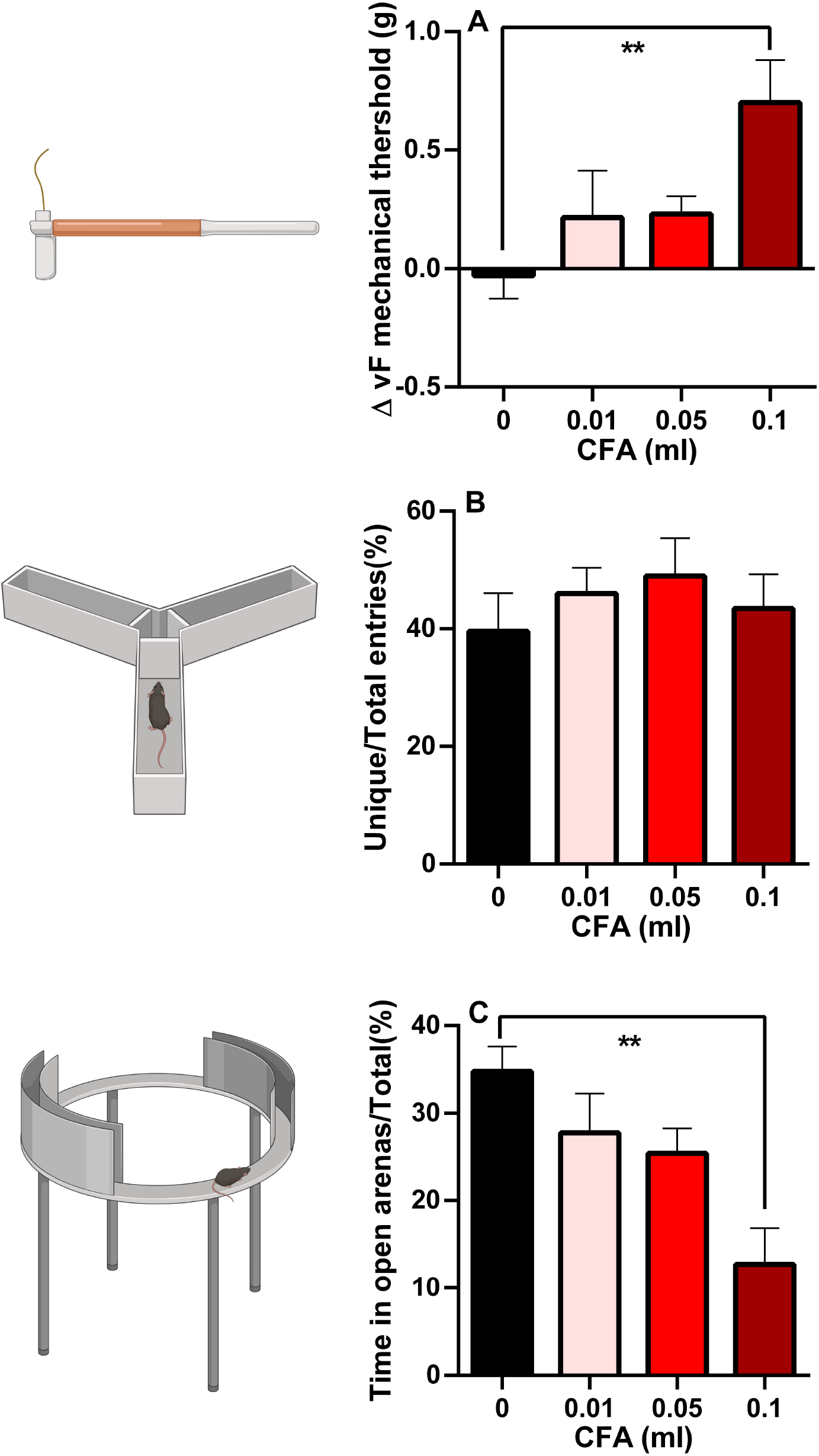
Visual illustration of the study paradigm demonstrating the control (saline) and the 3 experimental groups in the von Frey, y-maze, and zero-maze assays. **A**: Animals receiving 0.1 ml of CFA demonstrated reduced mechanical sensitivity thresholds on the affected hindpaw (ΔvF threshold (g) = threshold_contra_ (g)-threshold_ipsi_ (g)]. **B**: No differences were observed in the % of unique entries in the zero maze. **C**: Animals receiving 0.1 ml of CFA spent less time in the open arenas of the zero-maze. One-way ANOVA followed by Dunnett’s test for multiple comparisons; n=5/group; error bars indicate S.E.M. Figure created using BioRender©.

We did not detect any deficits in working memory using the y-maze test (**Figure 1B**; one-way ANOVA p=0.714, F=0.459). However, we detected signs of anxiety-like behaviors in the 0.1ml CFA group compared to control in the zero maze assay (**Figure 1C**; one-way ANOVA p=0.005, F=6.376; Dunnett’s multiple comparison test with an adjusted P value of 0.002, mean difference=22.07). Surprisingly, anxiety-like behavior was not detected in the open field assay, as measured by % time spend in the central 10% area (**Figure 2A, B;** one-way ANOVA p=0.870, F=0.236) or central 25% area (one-way ANOVA p=0.909, F=0.179).

**Figure 2:**
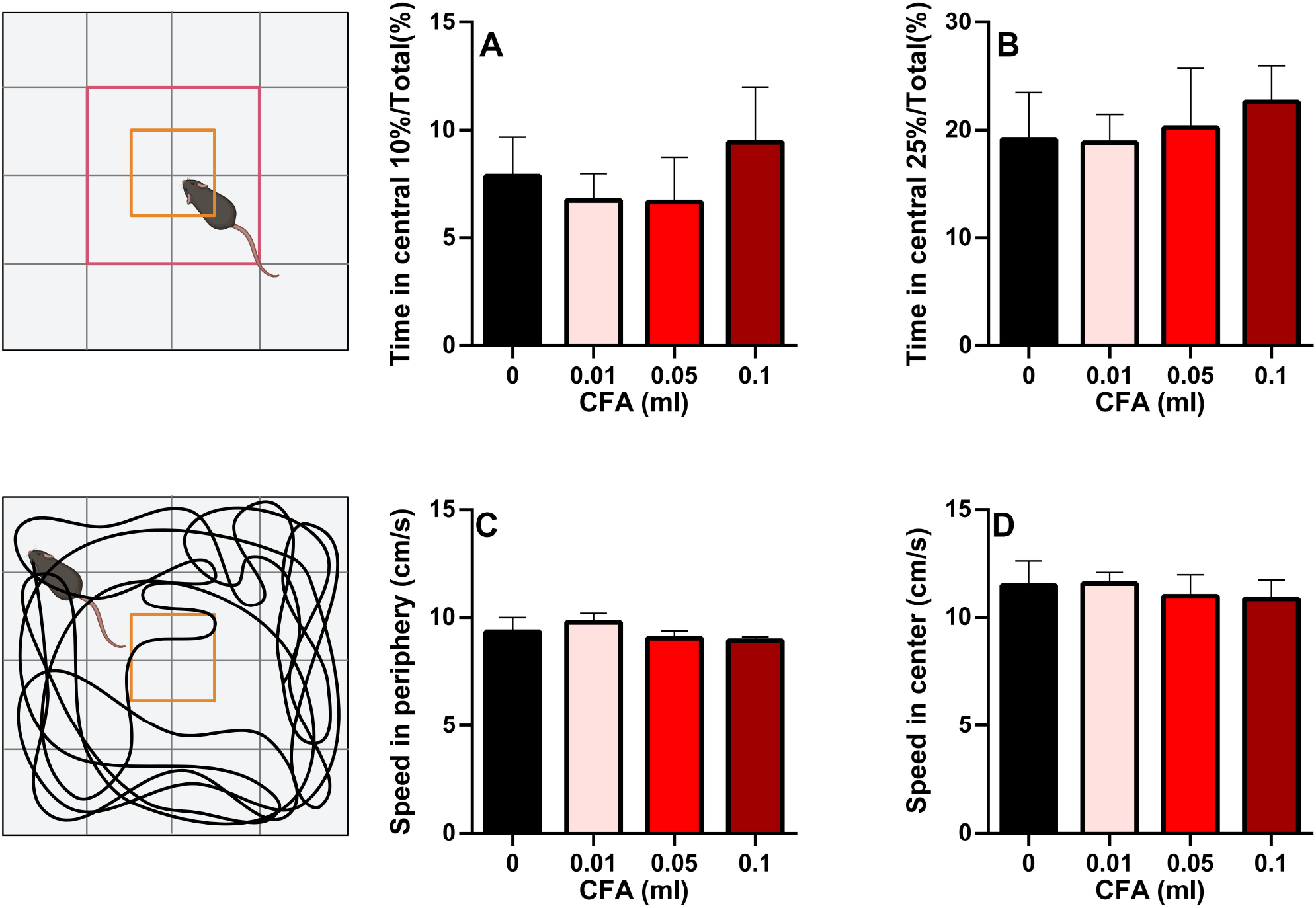
Visual illustration of the study paradigm demonstrating the control (saline) and the 3 experimental groups in the open field assay. **A, B**: No differences were observed in the time spent in the central 10% (A) or 25% (B) arena. **C, D**: No differences were observed in the speed of locomotion in the peripheral (C) or central (D) arenas of the apparatus (one-way ANOVA). n=5/group; error bars indicate S.E.M. Figure created using BioRender©.

Since many behavioral assays rely on motor function, we measured the speed of locomotion in the central and peripheral areas of the open field. No differences were observed in any of the groups (**Figure 2C, D;** peripheral area: one-way ANOVA p=0.144, F=2.071; central area: one-way ANOVA p=0.745, F=0.414)

Peripheral inflammation was associated with cerebral hypoperfusion in the parietal-temporal lobes in a dose-dependent manner 3 days after CFA administration [**Figure 3;** one-way ANOVA, p=0.0004, F (3, 20) = 9.55]. Compared to the saline control, 0.05 ml and 0.1ml CFA-treated groups demonstrated reduced perfusion rates (mean difference= 96.19 and 109.3, respectively).

**Figure 3:**
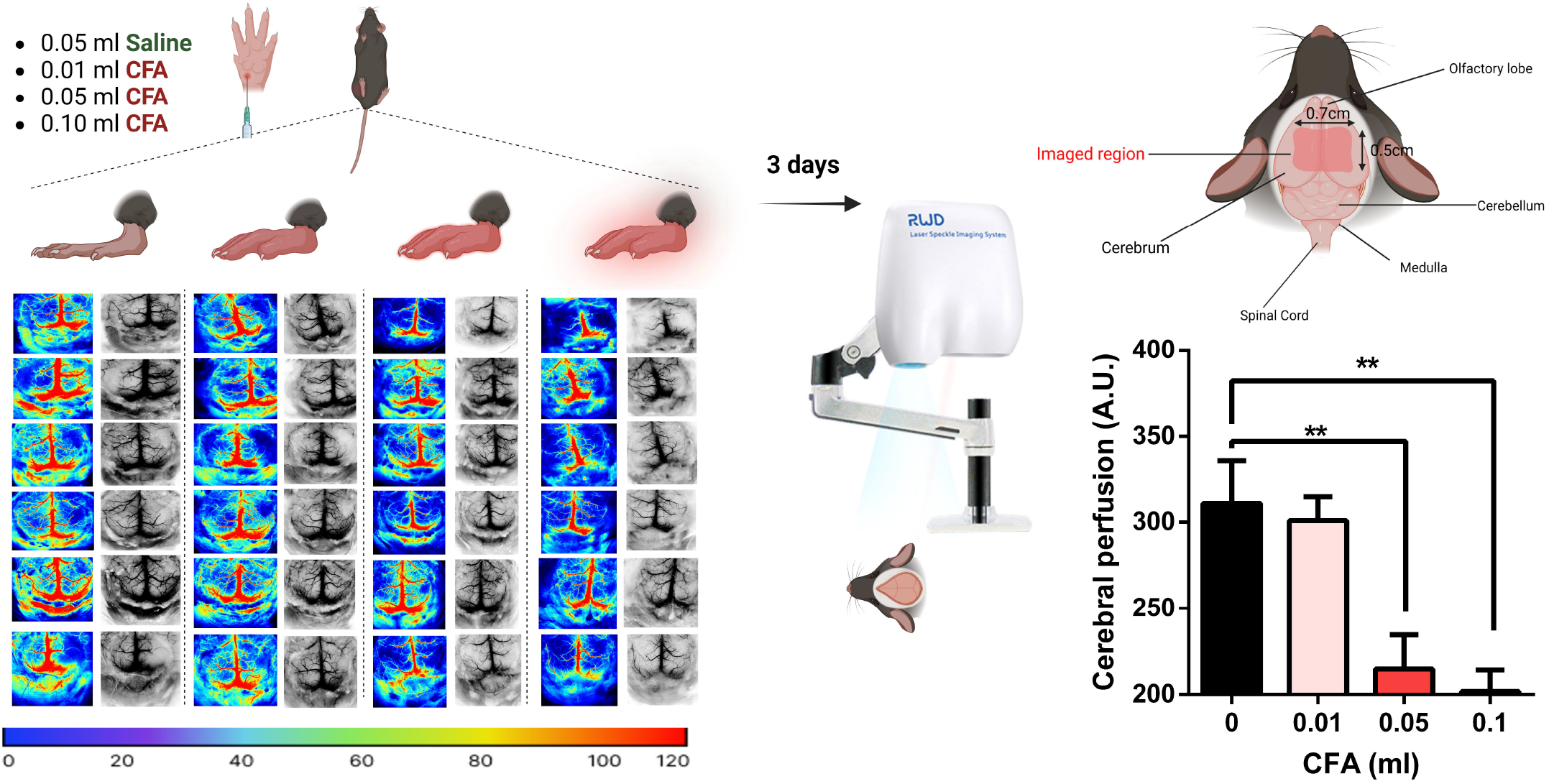
Visual illustration of the study paradigm demonstrating the control (saline) and the 3 experimental groups with the respective P-color images and gray cerebral blood flow patterns in twenty-four mice brains 3 days post-CFA/saline injection. Groups receiving 0.05 ml or 0.1 ml of CFA demonstrated reduced levels of cerebral perfusion (one-way ANOVA followed by Dunnett’s test for multiple comparisons). n=6/group; error bars indicate S.E.M. Figure created using BioRender©.

## Discussion

The CFA model of inflammation, while limited in its duration and severity of symptoms, is valuable in demonstrating the link between peripheral mechanisms of painful injury and the ensuing behavioral changes and biochemical and molecular alterations in central nervous system tissues. For example, hindpaw inflammation (0.02ml CFA) has been associated with stimulus-evoked hypersensitivity as well as measures of voluntary behavior alterations up to 9 days post-injection [25]. Additionally, CFA (0.05ml) administration in the rat hindpaw resulted in increased anxiety, increased levels of circulating corticosterone as well as decreased global DNA methylation levels in the amygdala 10 days after treatment [28]. In a separate study using in the same model, signs of increased blood-brain-barrier (BBB) permeability were paralleled by alterations in transmembrane tight junction protein levels [8]. Our results complement these observations by showing CFA-associated pain and anxiety as well as dose-dependent cerebral hypoperfusion 3 days following the onset of inflammation.

These findings can be viewed within the wider scope of peripheral inflammation using different models. For example, the murine lipopolysaccharide (LPS) model has repeatedly been associated with cognitive dysfunction [27] as well as aberrant synaptic phagocytosis by microglia [20], even prompting the hypothesis that LPS exposure could be a contributing factor to neurodegenerative disorders such as Alzheimer’s disease [9]. Inflammation subsequent to systemic LPS has also shown to be accompanied by ultrastructural cyto-architechtural changes in the BBB [16]. Finally, reduced perivascular flow of cerebrospinal fluid, in the absence of changes in blood flow, was observed in a murine LPS model [21]. There is some evidence for clinical translation as well: in patients with chronic neck and upper body pain, neck disability indices predicted cerebral hypoperfusion measured by single-photon emission computed tomography, with higher indices being linked to greater hypoperfusion [4]. In the more extreme case of sepsis, behavioral changes such as delirium and cognitive decline are linked to changes in cerebral microcirculation. For instance, both perfused cerebral vessel density and perfused small vessels were decreased in a ovine model of septic shock due to peritonitis [29].

We can therefore hypothesize that alterations in cerebral perfusion could present as a mechanistic link between peripheral inflammation and brain plasticity, and there is evidence for bidirectional modulation between the two. In models of chronic cerebral hypoperfusion, the extent of hypoperfusion is correlated to cognitive deficits [36] and vascular dementia is observed along with loss of BBB integrity, with a recent study demonstrating a role for the mechanosensitive piezo1 channel [34].

The adaptive value of cerebral hypoperfusion after peripheral inflammation remains uncertain. It is possible that hypoperfusion may aid in the minimization of inflammogen entry to the brain. It is also possible that it is secondary/compensatory instrument to systemic metabolic change since systemic inflammation in mice (LPS model) is associated with increased cerebral oxygen demand [19]. Future studies will target specific regions of interest, compare the right versus left hemispheres, and study timepoints beyond the 3-day window explored in this manuscript.

To our knowledge, this is the first study using laser speckle contrast imaging to examine the effects of CFA-induced peripheral inflammation on cerebral blood perfusion. The findings of decreased perfusion with increasing dose of CFA are useful in providing a mechanistic link between central and peripheral tissues in the processing of pain, and can provide insight into the underpinnings of centralization/chronification in other types of painful peripheral injuries.

## Declaration of Conflicting Interests

The authors declare no potential conflicts of interest with respect to the research, authorship, and/or publication of this article.

## Funding

Dr. Tajerian was supported by NIH grants 5SC2GM135114 and 1R16NS140306.

## Notes

### Competing Interest Statement

The authors have declared no competing interest.

## References

[1] Guide for the Care and Use of Laboratory Animals. Washington (DC), 2011.

[2] Alvarado S, Tajerian M, Millecamps M, Suderman M, Stone LS, Szyf M. Peripheral nerve injury is accompanied by chronic transcriptome-wide changes in the mouse prefrontal cortex. Mol Pain 2013;9:21.

[3] Alvarado S, Tajerian M, Suderman M, Machnes Z, Pierfelice S, Millecamps M, Stone LS, Szyf M. An epigenetic hypothesis for the genomic memory of pain. Front Cell Neurosci 2015;9:88.

[4] Bakhtadze MA, Vernon H, Karalkin AV, Pasha SP, Tomashevskiy IO, Soave D. Cerebral perfusion in patients with chronic neck and upper back pain: preliminary observations. J Manipulative Physiol Ther 2012;35(2):76–85.

[5] Becerra L, Morris S, Bazes S, Gostic R, Sherman S, Gostic J, Pendse G, Moulton E, Scrivani S, Keith D, Chizh B, Borsook D. Trigeminal neuropathic pain alters responses in CNS circuits to mechanical (brush) and thermal (cold and heat) stimuli. J Neurosci 2006;26(42):10646–10657.

[6] Berryman C, Stanton TR, Jane Bowering K, Tabor A, McFarlane A, Lorimer Moseley G. Evidence for working memory deficits in chronic pain: a systematic review and meta-analysis. Pain 2013;154(8):1181–1196.

[7] Borsook D, Hargreaves R. Brain imaging in migraine research. Headache 2010;50(9):1523–1527.

[8] Brooks TA, Hawkins BT, Huber JD, Egleton RD, Davis TP. Chronic inflammatory pain leads to increased blood-brain barrier permeability and tight junction protein alterations. Am J Physiol Heart Circ Physiol 2005;289(2):H738–743.

[9] Brown GC, Heneka MT. The endotoxin hypothesis of Alzheimer’s disease. Mol Neurodegener 2024;19(1):30.

[10] Cauda F, D’Agata F, Sacco K, Duca S, Cocito D, Paolasso I, Isoardo G, Geminiani G. Altered resting state attentional networks in diabetic neuropathic pain. J Neurol Neurosurg Psychiatry 2010;81(7):806–811.

[11] Cauda F, Sacco K, Duca S, Cocito D, D’Agata F, Geminiani GC, Canavero S. Altered resting state in diabetic neuropathic pain. PLoS One 2009;4(2):e4542.

[12] Chaplan SR, Bach FW, Pogrel JW, Chung JM, Yaksh TL. Quantitative assessment of tactile allodynia in the rat paw. J Neurosci Methods 1994;53(1):55–63.

[13] Cook AD, Christensen AD, Tewari D, McMahon SB, Hamilton JA. Immune Cytokines and Their Receptors in Inflammatory Pain. Trends Immunol 2018;39(3):240–255.

[14] DaSilva AF, Becerra L, Pendse G, Chizh B, Tully S, Borsook D. Colocalized structural and functional changes in the cortex of patients with trigeminal neuropathic pain. PLoS One 2008;3(10):e3396.

[15] Dimanche A, Miller DR, Goldberg J, Raabe A, Dunn AK, Bervini D. Continuous hemodynamics monitoring during arteriovenous malformation microsurgical resection with laser speckle contrast imaging: case report. Front Surg 2023;10:1285758.

[16] Erickson MA, Shulyatnikova T, Banks WA, Hayden MR. Ultrastructural Remodeling of the Blood-Brain Barrier and Neurovascular Unit by Lipopolysaccharide-Induced Neuroinflammation. Int J Mol Sci 2023;24(2).

[17] Geha PY, Baliki MN, Harden RN, Bauer WR, Parrish TB, Apkarian AV. The brain in chronic CRPS pain: abnormal gray-white matter interactions in emotional and autonomic regions. Neuron 2008;60(4):570–581.

[18] Harris RE, Sundgren PC, Craig AD, Kirshenbaum E, Sen A, Napadow V, Clauw DJ. Elevated insular glutamate in fibromyalgia is associated with experimental pain. Arthritis Rheum 2009;60(10):3146–3152.

[19] Liu C, Cardenas-Rivera A, Teitelbaum S, Birmingham A, Alfadhel M, Yaseen MA. Neuroinflammation increases oxygen extraction in a mouse model of Alzheimer’s disease. Alzheimers Res Ther 2024;16(1):78.

[20] Manabe T, Racz I, Schwartz S, Oberle L, Santarelli F, Emmrich JV, Neher JJ, Heneka MT. Systemic inflammation induced the delayed reduction of excitatory synapses in the CA3 during ageing. J Neurochem 2021;159(3):525–542.

[21] Manouchehrian O, Ramos M, Bachiller S, Lundgaard I, Deierborg T. Acute systemic LPS-exposure impairs perivascular CSF distribution in mice. J Neuroinflammation 2021;18(1):34.

[22] McWilliams LA, Goodwin RD, Cox BJ. Depression and anxiety associated with three pain conditions: results from a nationally representative sample. Pain 2004;111(1-2):77-83.

[23] Napadow V, Kim J, Clauw DJ, Harris RE. Decreased intrinsic brain connectivity is associated with reduced clinical pain in fibromyalgia. Arthritis Rheum 2012;64(7):2398–2403.

[24] Parthasarathy AB, Weber EL, Richards LM, Fox DJ, Dunn AK. Laser speckle contrast imaging of cerebral blood flow in humans during neurosurgery: a pilot clinical study. J Biomed Opt 2010;15(6):066030.

[25] Pitzer C, Kuner R, Tappe-Theodor A. Voluntary and evoked behavioral correlates in inflammatory pain conditions under different social housing conditions. Pain Rep 2016;1(1):e564.

[26] Seifert F, Kiefer G, DeCol R, Schmelz M, Maihofner C. Differential endogenous pain modulation in complex-regional pain syndrome. Brain 2009;132(Pt 3):788–800.

[27] Shaw KN, Commins S, O’Mara SM. Lipopolysaccharide causes deficits in spatial learning in the watermaze but not in BDNF expression in the rat dentate gyrus. Behav Brain Res 2001;124(1):47–54.

[28] Spinieli RL, Cazuza RA, Sales AJ, Carolino ROG, Martinez D, Anselmo-Franci J, Tajerian M, Leite-Panissi CR. Persistent inflammatory pain is linked with anxiety-like behaviors, increased blood corticosterone, and reduced global DNA methylation in the rat amygdala. Mol Pain 2022;18:17448069221121307.

[29] Taccone FS, Su F, Pierrakos C, He X, James S, Dewitte O, Vincent JL, De Backer D. Cerebral microcirculation is impaired during sepsis: an experimental study. Crit Care 2010;14(4):R140.

[30] Tajerian M, Alvarado S, Millecamps M, Vachon P, Crosby C, Bushnell MC, Szyf M, Stone LS. Peripheral nerve injury is associated with chronic, reversible changes in global DNA methylation in the mouse prefrontal cortex. PLoS One 2013;8(1):e55259.

[31] Tajerian M, Alvarado SG, Clark JD. Differential olfactory bulb methylation and hydroxymethylation are linked to odor location memory bias in injured mice. Mol Pain 2019;15:1744806919873475.

[32] Tajerian M, Hung V, Nguyen H, Lee G, Joubert LM, Malkovskiy AV, Zou B, Xie S, Huang TT, Clark JD. The hippocampal extracellular matrix regulates pain and memory after injury. Mol Psychiatry 2018;23(12):2302–2313.

[33] Tajerian M, Leu D, Zou Y, Sahbaie P, Li W, Khan H, Hsu V, Kingery W, Huang TT, Becerra L, Clark JD. Brain neuroplastic changes accompany anxiety and memory deficits in a model of complex regional pain syndrome. Anesthesiology 2014;121(4):852–865.

[34] Xu F, Xin Q, Ren M, Shi P, Wang B. Inhibition of piezo1 prevents chronic cerebral hypoperfusion-induced cognitive impairment and blood brain barrier disruption. Neurochem Int 2024;175:105702.

[35] Zhang D, Wang W, Zhu X, Li R, Liu W, Chen M, Vu T, Jiang L, Zhou Q, Evans CL, Turner DA, Sheng H, Levy JH, Luo J, Yang W, Yao J, Hoffmann U. Epinephrine-induced Effects on Cerebral Microcirculation and Oxygenation Dynamics Using Multimodal Monitoring and Functional Photoacoustic Microscopy. Anesthesiology 2023;139(2):173–185.

[36] Zhou Z, Ma Y, Xu T, Wu S, Yang GY, Ding J, Wang X. Deeper cerebral hypoperfusion leads to spatial cognitive impairment in mice. Stroke Vasc Neurol 2022;7(6):527–533.

